## Supplemental Figures for "Mixotrophy for carbon-conserving waste upcycling"

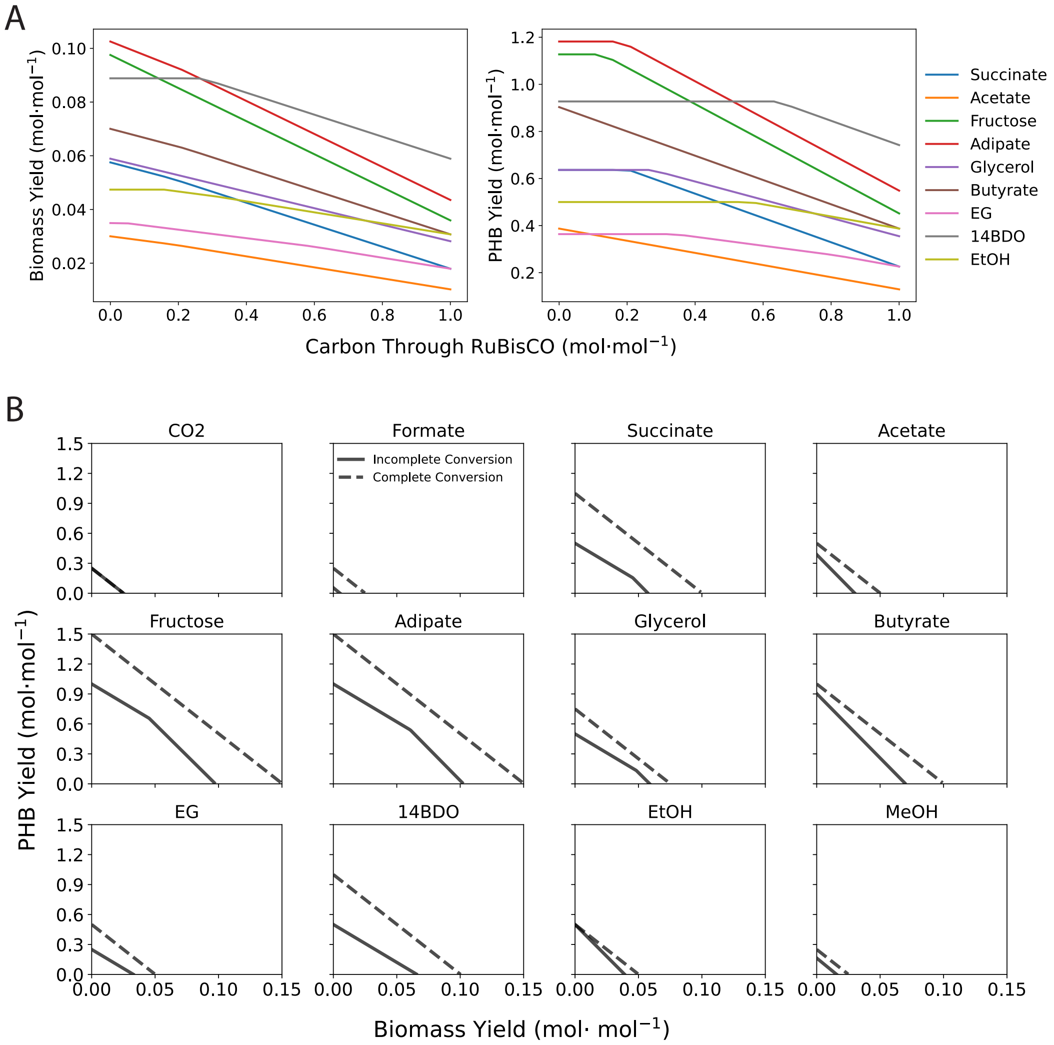


**Figure 1.** (A) Carbon reassimilation via RuBisCO without an additional electron source is detrimental to both biomass and PHB yield. A fraction of total carbon flux was forced through the RUBISCO reaction with $H_2$ uptake blocked and biomass or PHB production as the objective function. Yields decline as the proportion of carbon that passes through RUBISCO increases. (B) Complete carbon conversion with electrons from $H_2$ improves yields. For each carbon source, optimal PHB production for a given minimum biomass flux was determined, both with complete carbon conversion driven by $H_2$ oxidation and with $CO_2$ efflux allowed. Carbon reassimilation expands production envelopes for all carbon sources, yielding stoichiometric conversion to PHB (Fig. 1D).

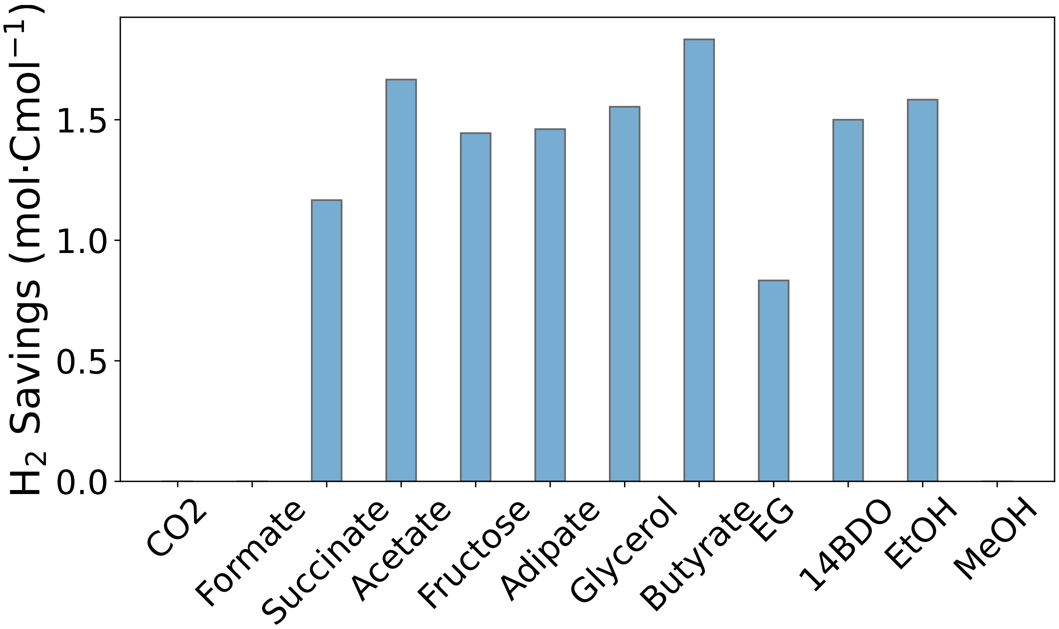


**Figure 2.** Hydrogen savings for PHB production with carbon conservation relative to the case in which all carbon is oxidized to carbon dioxide first, then reduced using hydrogen. C1 compounds must be assimilated this way, so there are no relative hydrogen savings for these feedstocks. Conversion of acetate, butyrate, and ethanol to PHB requires notably less hydrogen than in the full oxidation case, while ethylene glycol (EG) conversion involves minimal hydrogen savings.

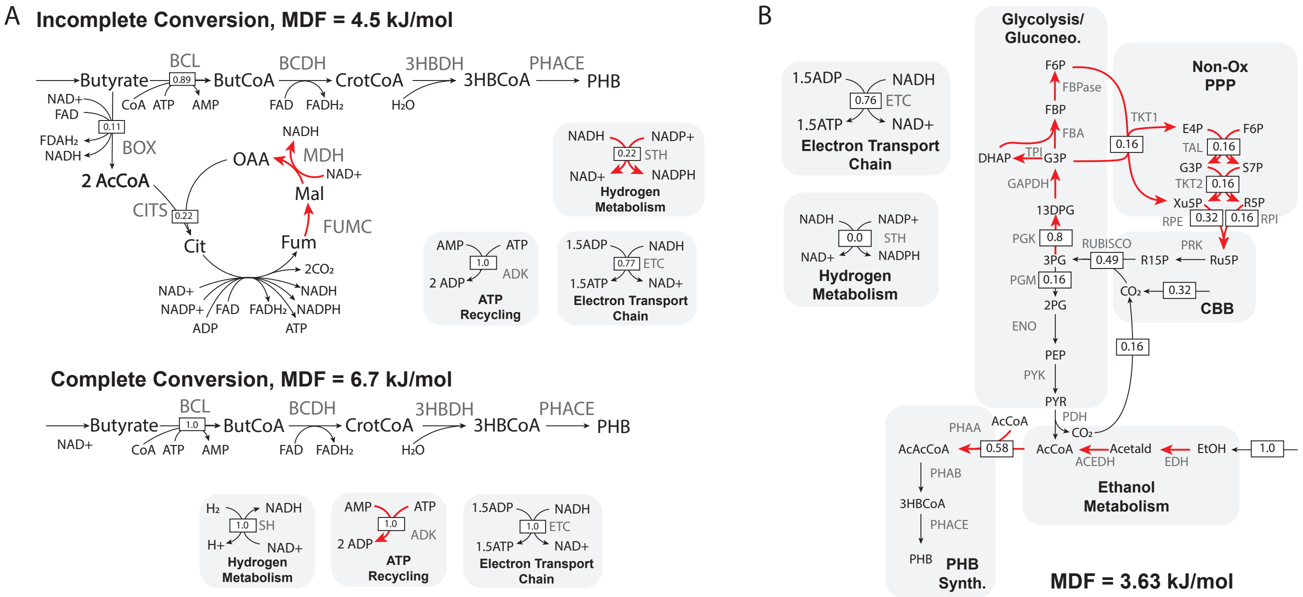


**Figure 3.** (A) Flux distribution for EtOH utilization. EtOH oxidation to acetyl-CoA yields sufficient reducing equivalents to drive polymerization and de novo carbon dioxide assimilation via the CBB. Each mole of EtOH used forces assimilation of 0.32 moles of carbon dioxide, yielding 0.58 moles of PHB. Reactions operating at the MDF of 3.63 kJ/mol are indicated in red. (B) Flux distributions for butyrate transformation to PHB with incomplete carbon conversion (top panel) and complete carbon conversion (bottom panel) with relative flux indicated. In the unforced scenario, 11% of assimilated butyrate must be converted to reducing equivalents via the carbon dioxide-producing TCA cycle. The requirement of fumarase ('FUMC') and malate dehydrogenase ('MDH') activities for energy generation yields an MDF of 4.5 kJ/mol. All reactions shown in red operate at this MDF. In the complete conversion scenario, hydrogen oxidation provides sufficient reducing power for all of the assimilated butyrate to be converted to PHB with no carbon dioxide generation, and therefore no reassimilation required. This reduces the thermodynamic constraint, allowing for operation at an MDF of 6.7 kJ/mol. EDH: ethanol dehydrogenase; ACEDH: acetaldehyde dehydrogenase; BCL: butyryl-CoA ligase; BCDH: butyryl-CoA dehydrogenase; 3HBDH: (R)-3-hydrobutyryl-CoA dehydratase; BOX: $\beta$-oxidation (lumped); Acetald: acetaldehyde; ButCoA: butyryl-CoA; CrotCoA: crotonyl-CoA

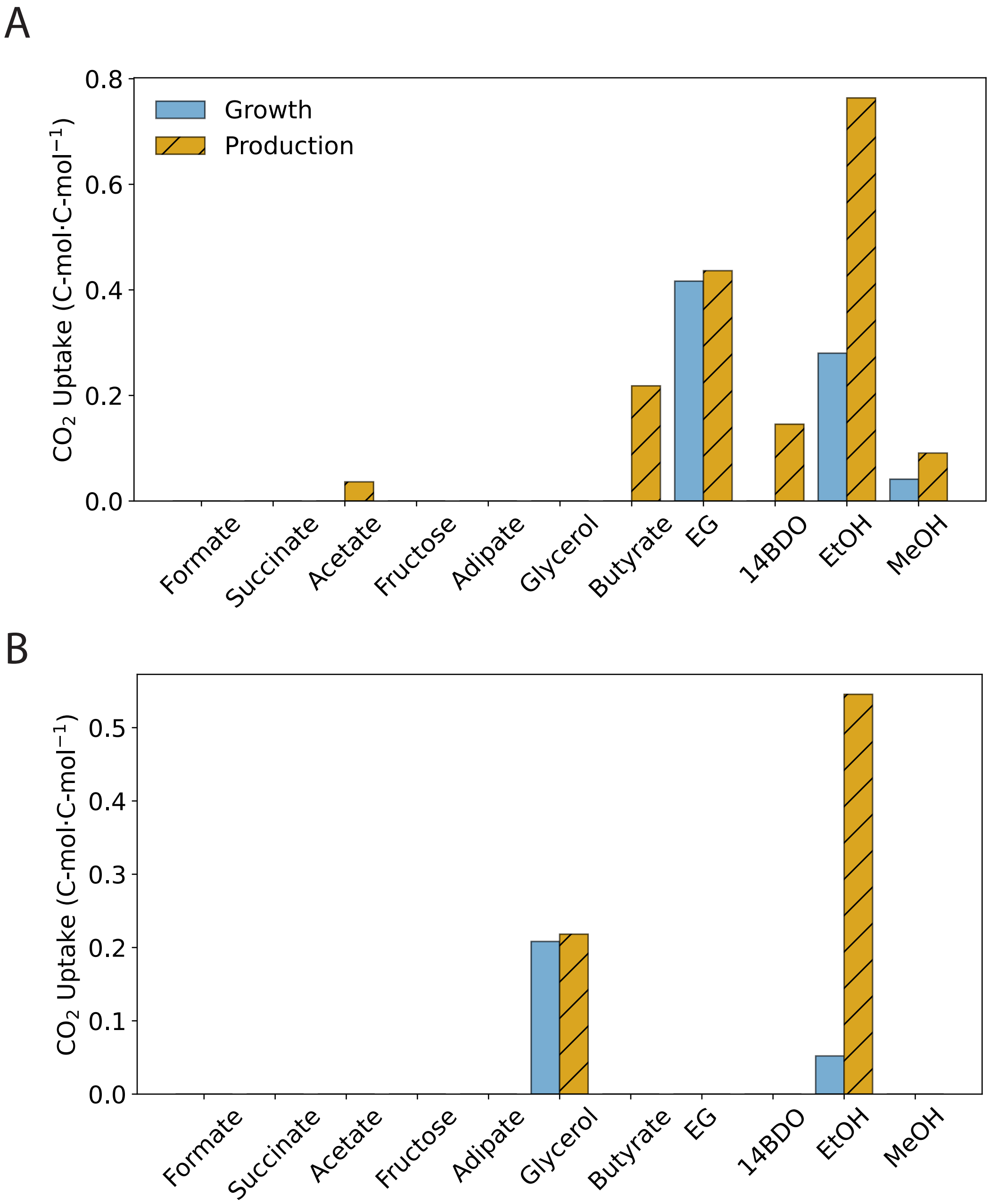


**Figure 4.** (A) De novo carbon dioxide assimilation is required for EG-to-GA mixotrophies. Here, EG is a source of electrons only, with carbon coming from the co-substrate and/or carbon dioxide. Partial oxidation of EG alone ('EG') requires carbon dioxide assimilation. (B) Glycerol-to-3HP mixotrophies do not require significant de novo carbon dioxide assimiliation. Partial oxidation of glycerol alone ('Glycerol') requires $CO_2$ assimilation.
